# Characterization of an operon required for growth on cellobiose in *Clostridioides difficile*

**DOI:** 10.1101/2021.06.26.450058

**Authors:** Md Kamrul Hasan, Babita Adhikari Dhungel, Revathi Govind

**Affiliations:** Division of Biology, Kansas State University, Manhattan, KS, 66506, USA

**Keywords:** *Clostridioides difficile*, *C. difficile*, *cel*R, cellobiose

## Abstract

Cellobiose metabolism is linked to the virulence properties in numerous bacterial pathogens. Here, we characterized a putative cellobiose PTS operon of *Clostridiodes difficile* to investigate the role of cellobiose metabolism in *C. difficile* pathogenesis. Our gene knockout experiments demonstrated that the putative cellobiose operon enables uptake of cellobiose into *C. difficile* and allows growth when cellobiose is provided as the sole carbon source in minimal medium. Additionally, using reporter gene fusion assays and DNA pull-down experiments, we show that its transcription is regulated by CelR, a novel transcriptional repressor protein, which directly binds to the upstream region of the cellobiose operon to control its expression. We have also identified cellobiose metabolism to play a significant role in *C. difficile* physiology as observed by the reduction of sporulation efficiency when cellobiose uptake was compromised in the mutant strain. In corroboration to *in vitro* study findings, our *in vivo* hamster challenge experiment showed a significant reduction of pathogenicity by the cellobiose mutant strain in both the primary and the recurrent infection model- substantiating the role of cellobiose metabolism in *C. difficile* pathogenesis.

## Introduction

Cellulose constitutes a significant portion of the human diet owing to its abundance in plant-based foods and vegetables. In the human digestive system, cellulose is broken down by microbial flora. In general, cellulose digestibility in humans is anywhere between 70-80% [1]. Although cellulose amounts for around 20% of the total carbohydrate consumed by the average human, it is a significant source of nutrients to the colon microbiota [2]. Several studies have been conducted to identify the gut microbiota population responsible for cellulose degradation [3]. Anaerobic cellulose degraders, which account for 5–10% of all cellulose decomposition, uses two different mechanisms for cellulose decomposition. The first is the use of a cellulosome complex of cellulolytic enzymes, which are known to occur in the outer cell envelope of the bacterial cell [4]. The second is the system in which cellulose is attached to the outer membrane through adhesins, including fibro-slime proteins and possibly type IV pilins [5, 6]. Both of these mechanisms work outside of the bacteria, causing diffusion of its major byproducts in the colon. One of the major byproducts of cellulose is the repeating units of cellobiose disaccharides. Cellobiose then becomes available to other gut-living microbes, which can uptake and utilize it as an alternative energy source. Thus, cellulose degradation and the potential use of its byproducts such as cellobiose are of considerable significance to the physiology of many enteric bacteria.

*C. difficile* can use a variety of carbohydrates and amino acids as its nutrient source [7]. Availability of nutrients is also crucial for *C. difficile* pathogenesis and disease progression, as *C. difficile* virulence factor gene expression is quite sensitive to nutrient availability [8–10]. Recent studies have indicated that carbohydrate metabolism (specifically complex carbohydrate metabolism) is a significant factor in many pathogenic bacteria’s virulence. *C. difficile* genome has multiple PTS operons capable of sequestering a host of different carbohydrates [9]. Complex carbohydrates such as cellobiose have been demonstrated to regulate virulence factors in several pathogenic bacteria [11, 12]. For example, in *Listeria monocytogenes*, a human pathogen naturally found in soils and decaying food products, cellobiose has been found to reduce the pathogenicity of bacteria by reducing the major virulence factor Listeriolysin and Phosphatidylinositol-specific phospholipase C [13]. In *L. monocytogenes*, cellobiose strongly represses virulence gene expression by inhibiting PrfA, the virulence gene activator [14]. Interestingly another study demonstrated the presence of cellobiose PTS operon to be responsible for increased neuro-invasiveness in hypervirulent *L. monocytogenes* strains [15]. In *Streptococcus pneumoniae*, when a genomic island containing a putative cellobiose PTS was deleted, the strain become attenuated in a murine pneumonia and sepsis model [16].

A large portion of the human diet consists of cellulose which can be converted into cellobiose by the other gut bacteria and can provide a sizable pool of extra nutrients for *C. difficile* to utilize. The ability to utilize cellobiose could lead to changes in the nutrient uptake and metabolism pathways in *C. difficile* in a way that might result in significant changes in the virulence properties of the bacteria. This study characterized the Cellobiose PTS operon in *C. difficile* R20291 (GenBank: FN545816.1) and has determined its importance in pathogenesis. We have also identified the transcriptional regulator, CelR, responsible for negatively regulating the Cellobiose PTS operon expression.

## Materials and Methods

### Bacterial strains and growth conditions

*C. difficile* strains (Table S1) were grown anaerobically in TY agar (Tryptose-Yeast extract) or 70:30 medium as described previously [17]. Cefoxitin (Cef; 25 µg/ml), thiamphenicol (Thio; 15 µg ml^-1^), and lincomycin (Lin; 15 µg ml^-1^) were added to *C. difficile* cultures whenever necessary. *Escherichia coli* strains were grown in (LB) broth. *E. coli strain* S17-1 were used for conjugation and supplemented with ampicillin (100 µg ml^-1^) or chloramphenicol (25 µg ml^-1^) when indicated and cultured aerobically in LB broth.

### Construction of R20291::*celA* and R20291::*celR* mutant strains

The mutants were constructed in a *C. difficile* strain using a ClosTron gene knockout system [18]. The group II intron insertion site was selected using the Perutka algorithm, a Web-based design tool available at http://clostron.com. The designed retargeted introns were cloned into pMTL007-CE5, and the resulting plasmids, pMTL007-CE5::Cdi-*celA*- 141a and pMTL007-CE5::Cdi-*cd2781*, was transferred into *C. difficile* R20291 by conjugation as described previously . The selection of thiamphenicol-resistant transconjugants in 15 µg ml^-1^ lincomycin plates confers potential ltrB (Ll.ltrB) insertions within the target genes in the chromosome of *C. difficile* R20291. The presence of putative mutants was identified by PCR using gene-specific primers (Table S2) in combination with the EBS universal primers. The R20291::*celR* (CDR20291_2781) was complemented by introducing pRG381 plasmid (Table S1) harboring *celR* into the mutant strain via conjugation.

### Growth Comparison between the R20291, R20291::*celA* and R20291::*celR*

Optical density of overnight cultures was adjusted to 1.0 at A600 nm and 100 µl of these cultures were used to inoculate 10 ml TY medium. We grew the cultures for 24 hours or longer in the anaerobic chamber at 37°C and measured the cell density by taking A600 nm at every 4 hours. The experiments were conducted in three replicates. *C. difficile* strains were also grown in CDMM (Completely defined minimal medium) [19] supplemented with either cellobiose (10 mM) or glucose (20 mM) or lichenin (0.1%) or chitobiose (0.1%) or N-acetylgucosamine (20 mM) as a sole source of carbon.

### Measurement of toxins using ELISA

Cytosolic toxins from 16h old *C. difficile* cultures grown in TY medium were measured as described previously [17]. In brief, one ml of *C. difficile* culture was harvested and suspended in 200 μl of sterile PBS supplemented with PMSF (2mM), sonicated, and centrifuged to harvest the cytosolic protein. One hundred micrograms of cytosolic proteins were used to measure the relative toxin levels using *C. difficile* premier Toxin A & B ELISA kit from Meridian Diagnostics Inc. (Cincinnati, OH). Toxin ELISA was performed in 3 replicates and independently repeated at least three times.

### Sporulation assay

Sporulation assay was performed as described previously. Briefly, C. difficile strains were grown on TY medium or TY supplemented with cellobiose (10 mM). After 30 h of growth, cells were scraped from the plates and suspended in TY to an OD600 of 1.0. To enumerate the viable vegetative cells and spores, cells were serially diluted and plated on to TY agar with 0.1% taurocholate and incubated at 37°C for 24 to 48 hours. To enumerate the number of viable spores only, 500 μl of the samples from each culture were mixed in a 1:1 ration with 95% ethanol and incubated for 30 minutes at room temperature to kill vegetative cells. The ethanol-treated samples were then serially diluted, plated on TY agar with 0.1% taurocholate and incubated at 37°C for 24 to 48 hours. The percentage of ethanol-resistant spores was calculated by dividing the number of CFU from spores by the total number of CFU and multiplying the value by 100. The results were based on a minimum of three biological replicates.

### Reporter gene fusion assay

Approximately 600 bps of the upstream DNA region of *celA*, or 200 bp upstream of CDR20291_2781 genes, along with their potential ribosomal-binding sites (RBS), were PCR amplified using specific primers with *KpnI* and *SacI* recognition sequences (Table S2) using R20219 chromosomal DNA as a template. Plasmid pRPF185 carries a *gusA* gene for beta-glucuronidase under the tetracycline-inducible (*tet*) promoter [20]. Using *KpnI* and *SacI* digestion, we removed the *tet* promoter and replaced it with either *celA*, or *CDR20219_2781* upstream regions to create plasmids pRG382, and pRG383 respectively. Plasmids with reporter fusions were then introduced into R20291, R20291::*celR* and R20291::*celA* mutant through conjugation as described above. The transconjugants were then grown in TY medium in the presence of appropriate antibiotics overnight. Overnight cultures were used as inocula at a 1:100 dilution to start a new culture. Bacterial cultures were harvested at every 4 hours of growth until 24 hours, and the amount of β-glucuronidase activity was assessed as described elsewhere with minor modifications [21, 22]. Briefly, the cells were washed, suspended in 0.8 ml of Z buffer (60 mM Na2HPO4 · 7H2O [pH 7.0], 40 mM NaH2PO4 · H2O, 10 mM KCl, 1 mM MgSO4· 7H2O, 50 mM 2-mercaptoethanol), and lysed using an equal amount of silica beads by bead beater method. The enzyme reaction was started by the addition of 0.16 ml of 6 mM *p*- nitrophenyl β-D-glucuronide (Sigma) to the broken cells and stopped by the addition of 0.4 ml of 1.0 M NaCO3. The β-Glucuronidase activity was calculated as described earlier [21, 22].

### Biotin DNA pulldown assays

Biotin pulldown assays were carried out as described elsewhere. Briefly, the DNA upstream to cellobiose operon was PCR amplified using biotin-labeled primers (Table S2). After gel extraction, the biotinylated DNA fragment was coupled to immobilized Monomeric Avidin Resin (G Biosciences) in B/W Buffer [23]. P*spoIIAB* (upstream of *spoIIAB*) and beads alone were treated alongside test samples as negative controls. The DNA and the beads were incubated at room temperature for 30 min in a rotor. The bead-DNA complex was washed with TE Buffer to remove any unbound DNA. To prepare cell lysates, *C. difficile* R20291::*celR* strain producing CelR-FLAG was grown to late exponential phase (16 h) in 500 ml TY medium at pH 7.4. After washing with 1XPBS, the cells were resuspended in BS/THES buffer and lysed using a French press. The whole lysate was centrifuged at 20,000 g for 30 min at 4°C. Lysate prepared from R20291::*celR* with vector alone control was also processed similarly and served as a control. The supernatants, along with salmon sperm DNA as a nonspecific competitor, was incubated with the bead-DNA complex and allowed to rotate at 4°C overnight. The bead-DNA-protein complex was washed with BS/THES Buffer (5 times). Elution was carried out with 50mM, 100 mM, and 200 mM NaCl in Tris-HCl pH 7.4. The eluates were analyzed by SDS-PAGE and western blotting using FLAG-tag specific antibody (GeneScript).

### Hamster model for *C. difficile* pathogenesis

Male and female Syrian Golden hamsters (100–120 g) were used for *C. difficile* infection. Upon their arrival, fecal pellets were collected from all hamsters, homogenized in 1 ml saline, and examined for *C. difficile* by plating on CCFA-TA (Cycloserine Cefoxitin Fructose Agar- 0.1% Taurocholate) to ensure that the animals did not harbor indigenous *C. difficile*. After this initial screen, they were housed individually in sterile cages with ad libitum access to food and water for the duration of the study. Hamsters were first gavaged with 30 mg kg^-1^ clindamycin [24, 25]. *C. difficile* infection was initiated five days after clindamycin administration by gavage with vegetative cells. We used vegetative *C. difficile* cells because the test strain (R20291::*celA*) produces very few spores. Bacterial inoculums were standardized and prepared immediately before challenge as described in our earlier study [25]. They were transported in independent 1.5 ml Eppendorf tubes to the vivarium using the Remel AnaeroPack system (one box for each strain) to maintain viability. Immediately before and after infecting the animal, a 10 μL sample of the inoculum was plated onto TY agar with cefoxitin to confirm the bacterial count and viability. There were three groups of animals, including the uninfected control group. Ten animals per group were used for the infection. Approximately, 1500 *C. difficile* vegetative cells of R20291 strain and R20291::*celA* were used for the animal challenge. In the uninfected control (group 3), only six animals were used, and they received only antibiotics and sterile PBS. Hamsters were monitored for signs of disease (lethargy, poor fur coat, sunken eyes, hunched posture, and wet tail) every four hours (six times per day) throughout the study period. Hamsters were scored from 1 to 5 for the signs mentioned above (1-normal and 5-severe). Hamsters showing signs of severe disease (a cumulative score of 12 or above) were euthanized by CO2 asphyxiation.

For the recurrent infection model study, hamsters recovered from the primary treatment were used. Three weeks after the recovery, animals were treated with Clindamycin without further challenge with *C. difficile*. Surviving hamsters were euthanized 10 days later. The cecal contents harvested from the sacrificed hamsters were collected in 15ml Nalgene tubes, secured airtight, and transported to the lab using the Remel AnaeroPack system. They were then immediately subjected to CFU enumeration as described previously. The survival data of the challenged animals were graphed as Kaplan-Meier survival analyses and compared for statistical significance using the log-rank test using GraphPad Prism 7 software (GraphPad Software, San Diego, CA).

### Ethics statement

All animal procedures were performed with prior approval from the KSU Institutional Animal Care and Use Committee (protocol #4161). Animals showing signs of disease were euthanized by CO^2^ asphyxia followed by thoracotomy as a secondary means of death, in accordance with Panel on Euthanasia of the American Veterinary Medical Association. Kansas State University is accredited by AAALAC International (Unit #000667) and files an Assurance Statement with the NIH Office of Laboratory Animal Welfare (OLAW). KSU Animal Welfare Assurance Number is D16-00369 (A3609-01), and USDA Certificate Number is 48-R-0001. Kansas State University utilizes the United States Government Principles for the utilization and care of vertebrate animals used in testing, research, and training guidelines for appropriate animal use in a research and teaching setting.

## Results

### Cellobiose operon in *C. difficile*

The *C. difficile* R20291 strain harbors a putative cellobiose operon mapping from position 3287614 to 3291734 in its genome (GenBank: FN545816.1) (Fig. 1A). The operon contains five genes and is similar in genomic content to many other bacteria-harboring cellobiose transport and utilization operon, including *Bacillus subtilis* (Fig. S1) [26]. The operon consists of the genes *celA, celB, celF, CDR20291_2777*, and *celC*, in the respective order of arrangement (Fig. 1A). The operon encodes the multidomain phosphoenolpyruvate-dependent sugar-phosphotransferase (PTS) components EIIA, EIIB, and EIIC by *celC, celA*, and *celB*, respectively (Fig. 1B) [27]. The PTS components are capable of concurrent import and phosphorylation of cellobiose to cellobiose-6P. The *celF* encodes a 6-phospho beta-glucosidase, which catalyzes the breakdown of phosphorylated cellobiose-6P to Glucose and 6-phospho glucose [28]. BlastP identifies CDR20291_2777 as a putative chitin disaccharide deacetylase, an enzyme capable of producing heterodisaccharide from its substrate chitobiose [29].

**Figure 1.**
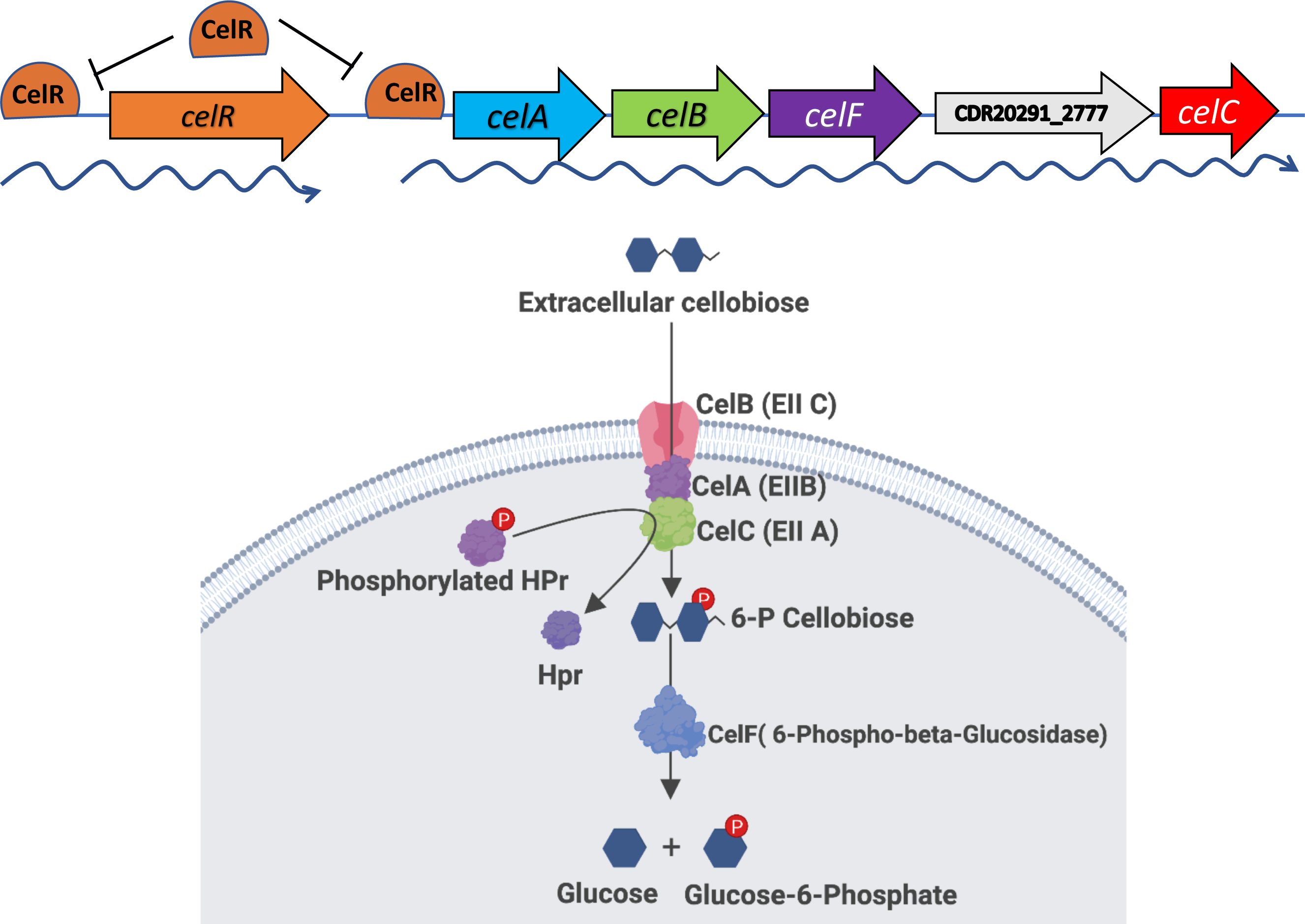
Schematics of Cellobiose operon and PTS system. **A.** The Cellobiose PTS operon of the *C. difficile* R20291 genome consists of *celA, celB, celF, CDR20291_2777,* and *celC* (*cd630_2884, cd630_2883, cd630_2882, cd630_2881, and cd630_2880* respectively in *C. difficile* 630 strain). CelR, the repressor of the operon, represses its own expression as well. **B.** The PTS consists of two cytoplasmic energy-coupling proteins (Enzyme I and HPr) and carbohydrate-specific Enzymes II, which catalyze concomitant carbohydrate translocation and phosphorylation of the substrate. In *C. difficile* extracellular Cellobiose is potentially uptake by a PTS system consisting of CelB, CelA, and CelC polypeptides making up the transmembrane EII part of the PTS system. The phosphate group could be transferred to the PTS system by the HPr. Phosphorylated Cellobiose could then be broken down to Glucose and Glucose 6- Phosphate by the enzymatic activity of CelF and incorporated in metabolic pathways.

Immediately upstream of the putative cellobiose PTS operon is the gene *CDR20291_2781* which codes for a putative GntR type transcriptional regulator. GntR type transcriptional regulators are found in various bacteria and are well documented to play roles in a multitude of cellular processes [30–34], including carbohydrate metabolism [35, 36]. The GntR regulators carry an N-terminal DNA binding domain and a C-terminal substrate-binding domain [37]. Although the C-terminal domain doesn’t bind with DNA, substrate-bound to this domain creates steric hindrance in the N-terminal Helix -Turn-Helix DNA binding domain and thus regulates its transcriptional regulatory activity [38, 39]. In many of the described cellobiose PTS operons, the transcription of the operon is controlled by a transcriptional regulator coded by a gene in proximity to the operon [40–42]. It is also known that many other sugar PTS operons are under transcriptional regulation by GntR type regulators. Some examples include GmuR repressor for glucomannan utilization operon in *Bacillus subtilis* [43] and GntR regulator controlling Gluconate utilization operon in *Vibrio cholerae* [44]. In *Streptococcus pneumoniae,* the GntR type regulators CelR and BguR are demonstrated to regulate the expression of *cel* and *bgu* operons involved in cellobiose uptake and metabolism [40, 45]. Based on these examples, we hypothesized that CDR20291_2781 has a high potential to be a transcriptional regulator of the putative cellobiose PTS operon in *C. difficile*, which we subsequently tested.

### Cellobiose metabolism effects on *C. difficile* growth

To understand the importance of cellobiose utilization in *C. difficile* physiology and virulence, we disrupted *celA*, the first gene in the putative cellobiose operon, by inserting a group II intron using the Clostron technique [18]. PCR using intron-specific and gene-specific primers were performed to confirm the insertion (Fig. S2). RT-PCR and qRT-PCR were performed to check the effect of insertion on the transcription of downstream genes. No transcripts of *celB, celC, celF,* and *CD2777* were detected in the R20291::*celA* mutant showing that the disruption of the first gene of the operon had disrupted the transcription of the entire operon (Fig. S3). We disrupted the putative GntR regulator CD2781 (here forth called *celR*) using the same Clostron technique described above (Fig. S2).

We examined the impact of cellobiose operon disruption on the growth of *C. difficile* in rich TY medium. In TY, there was no significant difference in growth dynamics between the parent, the R20291::*celR*, and R20291::*celA* strains (Fig 2A). We then compared the growth of parent and mutant strains in a minimum media where either cellobiose, lichenin, glucose, chitibiose, or N-acetylglucosamine (GlcNAc) was provided as the sole carbon source. None of the *C. difficile* strains grew when lichenin was used as a sole carbon source, indicating the absence of a lichenin utilization mechanism in *C. difficile* R20291 strain (data not shown). Growth was observed in the parent strain and also in R20291::*celA* mutant in the presence of GlcNAc or chitobiose, indicating that the cellobiose operon is not involved in transportation and utilization of these sugars (Fig. S4).

**Figure 2.**
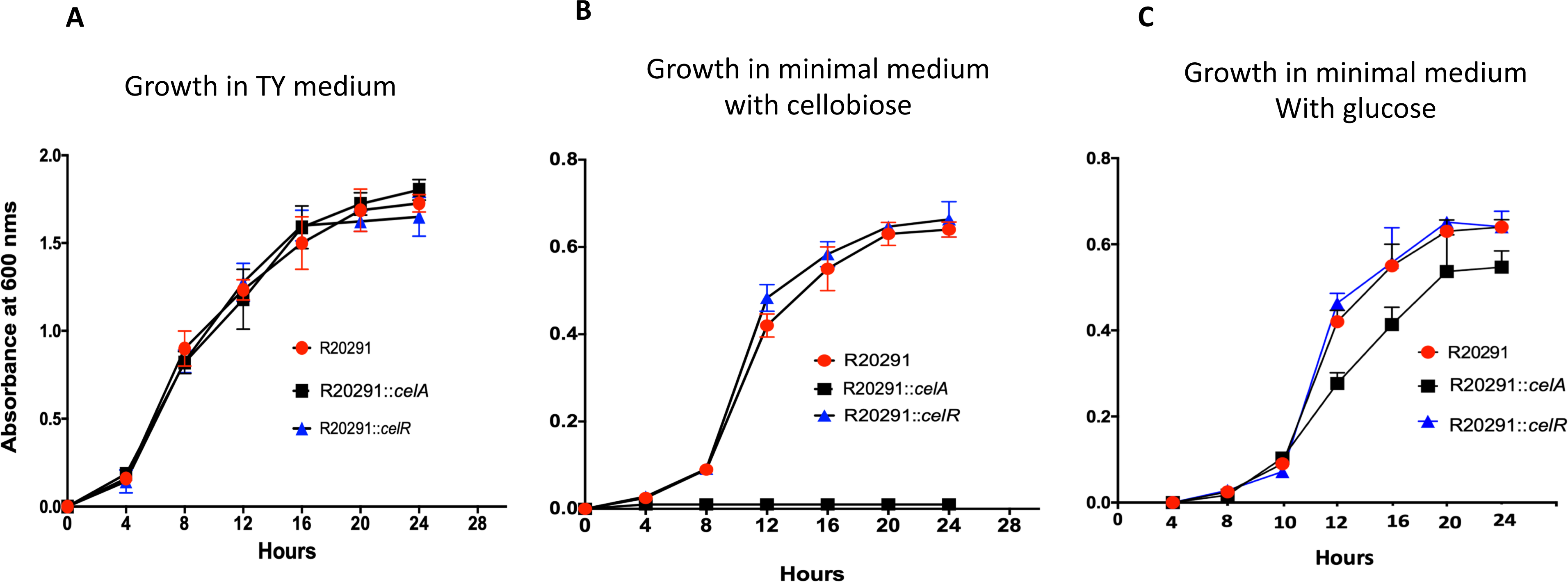
Role of cellobiose PTS operon on *C. difficile R20291* growth. **A.** Growth comparison between R20291, R20291::*celR*, and R20291::*celA* strains. Strains were grown in TY media and OD600 nm was measured every 4 hours and plotted. Growth comparison between R20291, R20291::*celR*, and R20291::*celA* strains grown in minimal media supplemented with 10 mM Cellobiose **(B)** or 20 mM glucose **(C).** Each experiment was repeated at least three times.

The *C. difficile* R20291::*celA* was unable to grow when cellobiose was provided as a sole carbon source, indicating the cellobiose PTS operon is responsible for the uptake and metabolism of cellobiose (Fig. 2B). *C. difficile* R20291::*celA* strain’s growth defect was partly restored in minimum media supplemented with glucose (Fig. 2C). This result suggests that cellobiose PTS system may also be partly responsible for transporting glucose.

This phenomena has been observed in other Gram-positive bacteria as well. For example, in several oral *streptococci*, glucose is transported by a PTS that also recognizes mannose, fructose, and the non-metabolizable analog 2-deoxyglucose [46, 47]. Growth defects were not observed in the R20291::*celR* strain in minimal media supplemented with either cellobiose, glucose, chitobiose, or GlcNAc (Fig. 2BC, Fig. S4).

### Cellobiose operon transcription is regulated by CelR, a GntR class regulator

To test our hypothesis that *celR* is a transcriptional regulator of the putative Cellobiose PTS operon, we created a promoter fusion in pRPF185 plasmid where the *tet* promoter of *gusA* gene coding for beta-glucuronidase was replaced by the 600 bp upstream region of cellobiose operon. The fusion plasmid was introduced in R20291 and R20291::*celR* strain to measure the beta-glucuronidase activity. Significantly increased level of reporter gene activity was observed in the mutant compared to parent strain indicating that transcription of the cellobiose operon in parent strain is repressed by CelR (Fig. 3A). We then conducted the beta-glucuronidase assay with the aforementioned strains grown in the presence of cellobiose or glucose. We observed a similar level of reporter gene activity between the parent and R20291::*celR* mutant strain (Fig. 3B) when grown in the presence of cellobiose, indicating that cellobiose could relieve CelR repression of the cellobiose operon.

**Figure 3.**
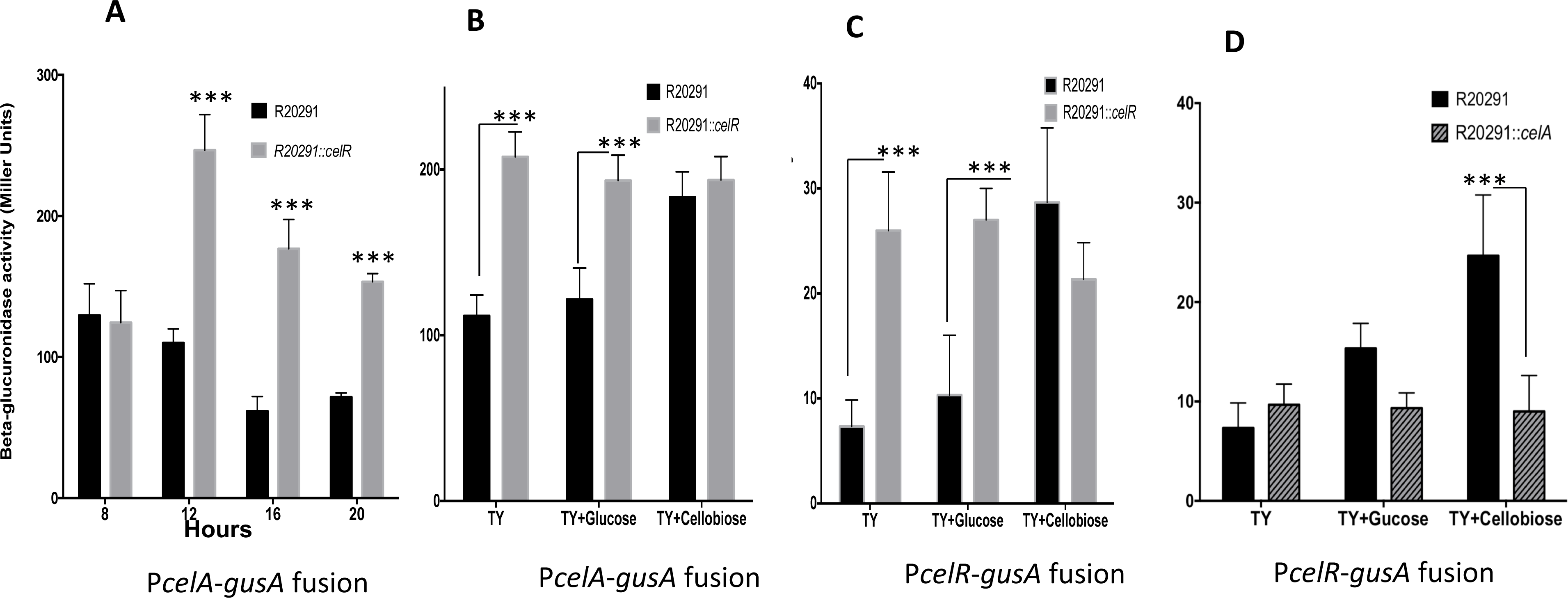
CelR mediated repression of Cellobiose PTS operon. **A.** Reporter assay with 600bp upstream region of *celA* in either R20291 parent or R20291::*celR* background. Strains were grown in TY and harvested at 8,12,16, and 20 hours timepoint and beta-glucuronidase activity was measured and plotted as Miller Units. Reporter assay of strains grown in TY or TY +glucose (20 mM) or TY+ cellobiose (10 mM): **(B)** Expression of *celA* promoter in R20291 and R20291::*celR* background, **(C)** Expression of *celR* promoter in R20291 and R20291::*celR* background **(D)** Expression of *celR* promoter in R20291 and R20291::*celA* background. Each experiment was repeated at least three times. Data were analyzed by Student’s two-tailed *t* test comparing parent strain R20291 with the mutants in the given growth condition. *** *p* value of <0.001; n=5.

Additionally, we found that CelR autoregulates its expression in a suppressive manner. We constructed a reporter fusion described in the earlier section with the promoter region of the *celR* gene and conducted a reporter activity assay. Results show a significant increase of activity in the *celR* mutant compared to the parent strain, indicating that CelR represses its transcription and the cellobiose operon (Fig. 3C). Similar to the CelR mediated repression of cellobiose operon, the autoregulation is relieved by the addition of cellobiose in the growth medium (Fig. 3C). We also measured the *celR* promoter activity in the R20291::*celA* strain background. We did not observe any change in the expression of *celR* promoter in the R20291::*celA* background even in the presence of cellobiose (Fig. 3D). This result suggests that processing of cellobiose is needed to regulate *celR* expression. Most likely, the CelR binding to cellobiose-P relieves the repression effect of CelR on its own promoter and the cellobiose operon.

Addition of glucose in the medium did not have any significant effect on the expression of the *cel* operon or the *celR* in the parent or in R20291::*celR*, indicating that cellobiose operon is not under catabolite repression (Fig. 3B, 3C). Interestingly, a slight increase in the expression of *celR* and *celA* promoter could be seen in the parent strain in the presence of glucose. This result again suggests that glucose might be transported using the components of cellobiose operon and might be partially relieving the CelR repression of *celR* and *cel* operon promoters. It is also important to note that transcription of *celR* and *cel* operon are present in the parent strain, even in the absence of cellobiose or glucose. One reason could be that other untested sugars in the TY medium could partially induce the expression of *celR* and the *cel* operon.

### CelR directly binds to the promoter region of Cellobiose PTS operon to repress transcription

The results in Fig. 3AB show that the reporter gene expression was less in the R20291 strain compared to R20291::*celR*, indicating CelR’s potential role as a repressor of the operon. To determine whether the repression of the cellobiose operon by CelR is due to its binding to the promoter region of the operon, we carried out a DNA binding experiment. Our initial attempts to carry out electro mobility shift assays (EMSA) using purified recombinant CelR-6His were unsuccessful (Data not shown). GntR regulators often need co-factors or undergo phosphorylation to bind to target [48]. To overcome this difficulty, we performed a biotin-labeled DNA pulldown assay to determine the DNA binding ability of CelR to the cellobiose operon’s promoter region under native conditions. The test DNA fragment was biotinylated and was coupled to immobilized Monomeric Avidin Resin. This bead-DNA complex was incubated with the cell lysate from the R20291::*celR* strain producing CelR-FLAG. The bound proteins were eluted, run in the SDS-PAGE, and immunoblotted with the FLAG-tag antibody. The biotinylated *spoIIBA* upstream DNA and the beads alone were also processed similarly and served as negative controls. Results showed that cellobiose operon upstream DNA could pulldown CelR-FLAG, suggesting that CelR binds specifically to the promoter region of the *lic* operon (Fig. 4AB). The DNA used for the pulldown contains an inverted repeat sequence separated by 23 bps. GntR regulators are known to bind of inverted repeat sequences. Secondary structure predictions of CelR suggest that it belongs to HutC family of GntR regulators [49]. This family of GntR regulators is known to bind to DNA sequence NyGT(A/C)TA(T/G)ACNy [31]. We could locate two of this repeats (AAAT**GTATATAC**ATTT) in the upstream of *celA* spanning the predicted promoter region (Fig 4A) [50]. To determine whether CelR binds to the repeat sequence, we synthesized a DNA fragment (Genewiz value genes) with mutations that potentially disrupt the consensus sequence and the inverted repeat and was used as a bait in the pulldown assay. CelR-FLAG failed to bind to the mutated DNA fragment, demonstrating the importance of this sequence in protein-DNA interaction. It is important to note that a similar consensus sequence (NyGT(A/C)TA(T/G)ACNy - AAAT**GTCTATAC**ATTTAA) is present upstream of *celR* gene as well, which may explain the autoregulation of *celR* we observed in the reporter assays.

**Figure 4.**
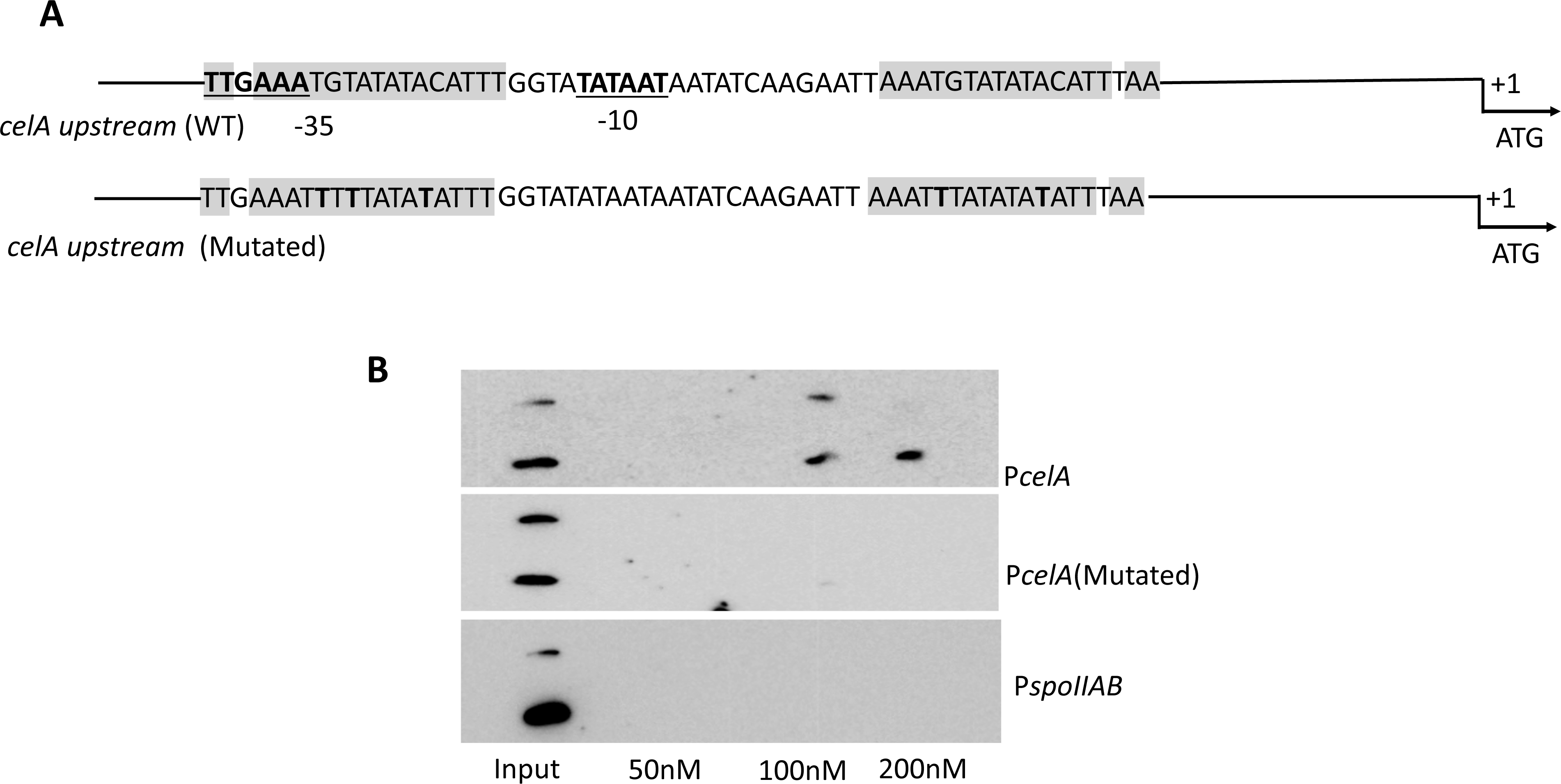
DNA pull down assay to determine CelR binding to the upstream DNA sequences of *celA*. **A.** DNA fragments used for the pull-down assay with the inverted repeat sequence highlighted and predicted promoter region (-10 and -35) underlined. P*celA* (mutated) DNA fragment carries point mutations in a potential CelR binding consensus sequence. **B.** Biotin labelled DNA pulldown assay with CelR. Cell lysate of strain expressing CelR- FLAG tagged was incubated with Biotin labelled immobilized test or control DNA fragments. Following wash, elution was done with 50mM, 100 mM, and 200 mM NaCl in Tris-HCl pH 7.4 and elutes were analyzed by Western Blot using anti-FLAG antibody.

### Cellobiose metabolism plays an imporatant role in *C. difficile* sporulation

A critical aspect of *C. difficile* pathogenesis is its capacity to produce toxins A and B and highly resilient exospores that can survive harsh physical and chemical treatment. To evaluate the role of cellobiose metabolism in *C. difficile* sporulation, we calculated the sporulation percentage of the R20291::*celA* and R20291::*celR* mutants in comparison to R20291 strain in TY medium with and without cellobiose. Our results show that R20291::*celA* mutant produces significantly fewer spores compared to the parent strain, in both growth conditions indicating *C. difficile* cellobiose metabolism’s role in sporulation (Fig. 5A). In contrast, the R20291::*celR* mutant produces a significantly higher number of spores compared to the parent strain (Fig. 5A, Fig. S5). We complemented the R20291::*celR* mutant by expressing CelR from a vector under its own promoter. We observed the CelR complemented strain to sporulate at a level comparable to the parent strain (Fig. 5A). In the presence of cellobiose, only the parent strain produced significantly less spores than in the absence of cellobiose. Presence of cellobiose did not influence the sporulation percentage in R20291::*celA* and R20291::*celR* mutant strains. In the *celR* complemented strain, however, a reduction can be seen when the strain was grown in the presence of cellobiose.

**Figure 5.**
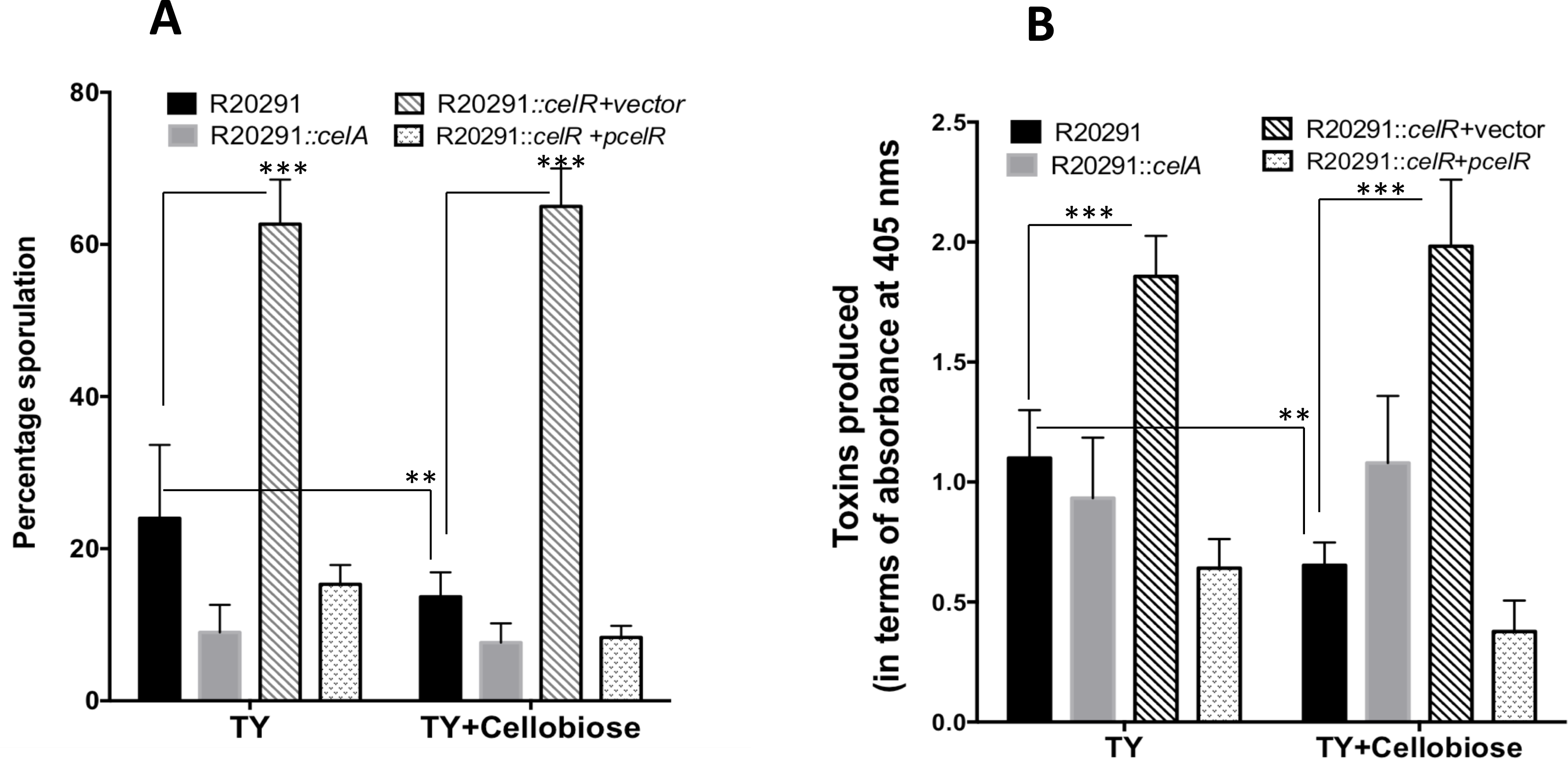
Role of cellobiose PTS operon on *C. difficile* R20291 sporulation and toxin production. **A.** Sporulation percentage of R20291, R20291::*celR*, R20291::*celA, and* R20291::*celR+pcelR* strains. Strains were grown in TY media or in TY supplemented with 10mM cellobiose. Sporulation percentage was calculated and plotted as described in the methods section**. B.** Toxin production level of R20291, R20291::*celR*, R20291::*celA, and* R20291::*celR+pcelR* strains. Strains were grown in TY or in TY+ cellobiose (10mM) up to 16 Hours and cytosolic toxin levels were measured using an ELISA based colorimetric assay and plotted as absorbance value @405nm. Each experiment was repeated at least three times. Data were analyzed by one-way ANOVA with Dunnett’s test for comparing R20291 with other strains used (multiple comparisons). ** Adjusted *p* value of <0.01; *** <0.001; n=4

To evaluate the role of cellobiose metabolism in *C. difficile* toxin production, we measured the cytosolic level of toxins of the R20291::*celA* and R20291::*celR* mutants and compared them with the R20291 parent strain. We observed no significant difference in toxin levels between R20291::*celA* mutant and parent strain (Fig. 5B). In contrast, the R20291::*celR* mutant produces a significantly higher amount of toxins than the parent strain (Fig. 5B). As expected, the CelR complemented strain produced toxins at a level comparable to the parent strain (Fig. 5B). Presence of cellobiose further reduced the toxin level in the *celR* complemented strain, suggesting a role for *celR* in toxin gene regulation.

### Cellobiose metabolism is essential for virulence and pathogenesis of recurrent CDI infection in hamsters

To determine the role of cellobiose metabolism in *C. difficile* pathogenesis, we challenged clindamycin-treated golden Syrian hamsters with approximately 1500 *C. difficile* R20291 and R20291::*celA* vegetative cells and monitored for infections for 10 days. We used vegetative cells since the R20291::*celA* strain sporulates poorly. Six animals out of ten in the parent group and one animal out of ten in the mutant group died by the end of the first challenge (Fig 6A). Bacterial load measurements detected high numbers of *C. difficile* in the cecal contents of the hamsters which died because of R20291 strain, but very few in the one hamster that died due to *R20291::celA* (Fig. 6C). This result suggests that the *R20291::celA* mutant is less capable of colonizing the host gut than the parent strain (*p* value <0.01, long rank test). The surviving hamsters were given clindamycin again to disrupt the gut microbiota and observed for the occurrence of a second round of infection, mimicking a recurrent infection in humans. None of the animals (None out of nine) infected with R20291::*celA* mutant strain had a recurrent infection, and *C. difficile* could not be recovered from their cecal contents. However, all of the animals (all four) infected with the parent strain got the recurrent infection and had to be euthanized before the end of the study period (Fig. 6B). Failure to cause reinfection can well be correlated with the poor sporulation capacity of the R20291::*celA* mutant. These results demonstrated that cellobiose PTS operon plays a vital role in *C. difficile*’s pathogenesis by affecting the sporulation and colonization of *C. difficile*.

**Figure 6.**
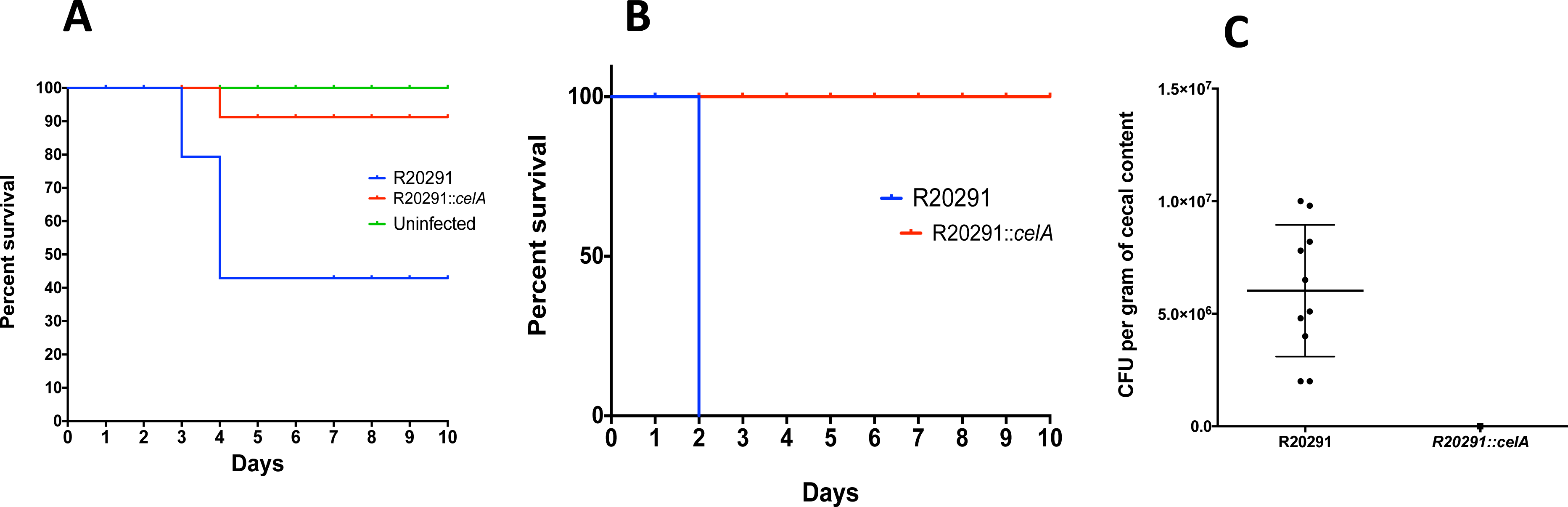
Role of cellobiose PTS operon on *C. difficile* R20291 pathogenicity in animal model. **A.** Kaplan-Meier survival curve of clindamycin-treated golden Syrian hamsters inoculated with *C. difficile* R20291 (n = 10) or R20291::*celA* mutant (n = 10). Six animals were used as an uninfected control. Animals were monitored every four hours for the symptoms of lethargy, poor fur coat, wet tail or hunched posture. Moribund animals were euthanized, and log-rank statistical analysis was performed. **B.** Kaplan-Meier survival curve of the Syrian golden hamsters in a recurrent infection study model. R20291 (n = 4) and R20291::*celA* mutant (n = 9) inoculated animals that survived the primary infection were retreated with clindamycin, leading to artificial induction of a recurrent infection. **C.** Comparison of bacterial load between R20291 parent and R20291::*celA* mutant strain infected animals. Cecal content collected from euthanized animals from primary infection was resuspended in 1X PBS, serially diluted and plated onto CCFA agar with 0.1% Taurocholate (CCFA-TA) to enumerate CFU counts.

## Discussion

A healthy human diet contains a large amount of plant-derived parts and vegetables that are rich in cellulose, which then get converted to cellobiose by different commensal bacteria. Although *C. difficile* primarily relies on the Stickland metabolism pathway for nutrient requirements [51], the capability to utilize alternative nutrient sources such as cellobiose abundant in the colon is undoubtedly beneficial. Toxin production and sporulation, the two main virulence determinants in *C. difficile*, are highly regulated by nutrient availability [9, 10, 52]. Hence understanding the cellobiose metabolism in *C. difficile* pathogenesis is important.

In this work, we identified a cellobiose PTS operon in *C. difficile*. We demonstrated that *C. difficile* was unable to utilize cellobiose when *celA*, the first gene of the operon, is disrupted. We also identified a novel GntR type transcriptional regulator- CelR of the Cellobiose PTS operon. The GntR type transcriptional regulators are known to positively and negatively regulate various bacteria’s carbohydrate metabolism pathways. Our growth study result shows that CelR is not essential for growth in cellobiose minimal media, ruling out its potential to be a positive regulator. Our reporter gene fusion assay demonstrates that CelR indeed works as a transcriptional repressor for the cellobiose operon. Although we saw that the addition of cellobiose relieves the repression of CelR on the cellobiose operon, we did not explore the exact mechanism of signal relay between the presence of cellobiose in the media to relieve repression of CelR. One hypothesis is that the cellobiose operon has a baseline expression allowing some cellobiose to be transported inside the bacteria, which could relieve the repression of CelR, hence increasing the expression of operon and uptake of more cellobiose. The CelR contains a UbiC transcription regulator-associated (UTRA) domain at C-terminal, which is known to modulate bacterial transcription factors in response to binding small molecules [31]. Hypothetically, cellobiose-6P could bind with the UTRA domain of CelR and induce conformational change that leads to disassociation from target DNA sequence, thus relieving repression.

Glucose is the preferred carbon source for many bacteria and represses the utilization of other sugars. In many Gram-positive bacteria, this carbon catabolite repression (CCR) is achieved by the global transcription regulator CcpA. In *C. difficile CcpA* is known to regulate differential fermentation pathways, selective utilization of carbon sources, amino acid metabolism, and toxin production in response to glucose availability [9]. Presence of glucose in the growth medium did not affect the expression of *cel* operon or its regulator *celR* (Fig. 3B, 3C), suggesting that its regulation is not under the regulation of CcpA. Transcriptome analysis of *ccpA* mutant in *C. difficile* failed to identify the *cel* operon as one of its targets [9]. Further, we could not identify the CcpA consensus binding site upstream of the *cel* operon or *celR*.

The R20291::*celA* strain showed a significant decrease in viable sporulation percentage, indicating cellobiose metabolism’s role in *C. difficile* physiology. When CelR repression of cellobiose operon is absent (in R20291::*celR* mutant strain), we observe an opposite phenotype of increased sporulation percentage. Our toxin assay data also shows that the R20291::*celR* produces a significantly higher amount of toxins than the parent strain. A similar phenomenon has been observed in a study describing a mutation in the trehalose uptake regulator TreR, causing increased virulence in *C. difficile* [8]. We observed an increased expression of *spo0A,* the sporulation master regulator and *tcdR,* the positive regulator of toxin genes in R20291::*celR* (Fig. S6). Since we identified the CelR binding consensus sequence, we used the motif search tool FIMO [53] to identify potential CelR binding regions in the R20291 genome. Interestingly, we identified a potential binding site upstream of CDR20291_1268 and CDR20291_1269, coding for a putative phosphodiesterase and its regulator, respectively (Table S3). Phosphodiesterase are enzymes that degrade the signaling molecule c-di-GMP. In *C. difficile*, c-di-GMP has been shown to negatively regulate toxin production and flagella mediated swimming motility [54–59].

We recently reported that phase variable expression of a phosphodiesterase (PdcB) could control sporulation in *C. difficile* [60]. Even though the mechanism of c-di-GMP mediated sporulation regulation is not known, we did report that reduced c-di-GMP concentration in *C. difficile* can increase the sporulation rate. We performed qRT-PCR to check the expression of CDR20291_1268 and CDR20291_1269 in the parent strain and in the *celR* mutant. We observed, nearly 7-fold and 1.5-fold increase in the expression of CDR20291_1268 and CDR20291_1269, respectively, in the *celR* mutant. This result suggests that CelR acts as a repressor and controls the production of CDR20291_1268 phosphodiesterase. Increased activity of the phosphodiesterase in the *celR* mutant would reduce the intracellular concentration of c-di-GMP and can positively influence the toxin production and sporulation in this mutant. However, an in-depth analysis is needed to prove this speculation.

Our animal model study demonstrates a functional cellobiose operon is important for *C. difficile* pathogenesis in hamsters. The R20291::*celA* mutant was significantly attenuated in the hamster both during initial and recurrent infection. We hypothesize that this phenomenon is a result of severely impaired spore production by R20291::*celA* mutant strain seen in our *in vitro* study. Indeed, our CFU count data shows a significant reduction in recovery of *C. difficile* from the R20291::*celA* challenged hamsters’ cecal content, indicating a rapid clearance of the bacteria from the host gut. Reduced pathogenicity could also result from R20291::*celA* mutant strain’s inability to utilize a vital carbon source such as cellobiose in a competitive gut microbiome, thus exhibiting a colonization disadvantage.

In summary, we have identified the PTS operon of *C. difficile*, which enables the bacteria to utilize cellobiose, a highly abundant alternate energy source, and demonstrated its implication in the bacteria’s pathogenicity. *C. difficile* infection and its associated complications are a major burden to the healthcare sector, and findings relating to its pathogenicity are of high significance. Our findings demonstrating cellobiose metabolism and its link to *C. difficile* virulence could help nutritionists provide a curated diet for vulnerable groups to minimize pathogenicity. Moreover, small molecule inhibitors or synthetic cellobiose analogues targeting the cellobiose PTS system could be designed and used as novel therapeutics to treat *C. difficile* infections in the future.

## Acknowledgement

We thank following investigators for sharing their lab resources: Nigel Minton, University of Nottingham, for the plasmid pMTL007C-E5; Robert Fagan for the vector pRPF185.

We thank Yusuf Ciftci for technical assistance throughout the study.

---

**Figure S1.**
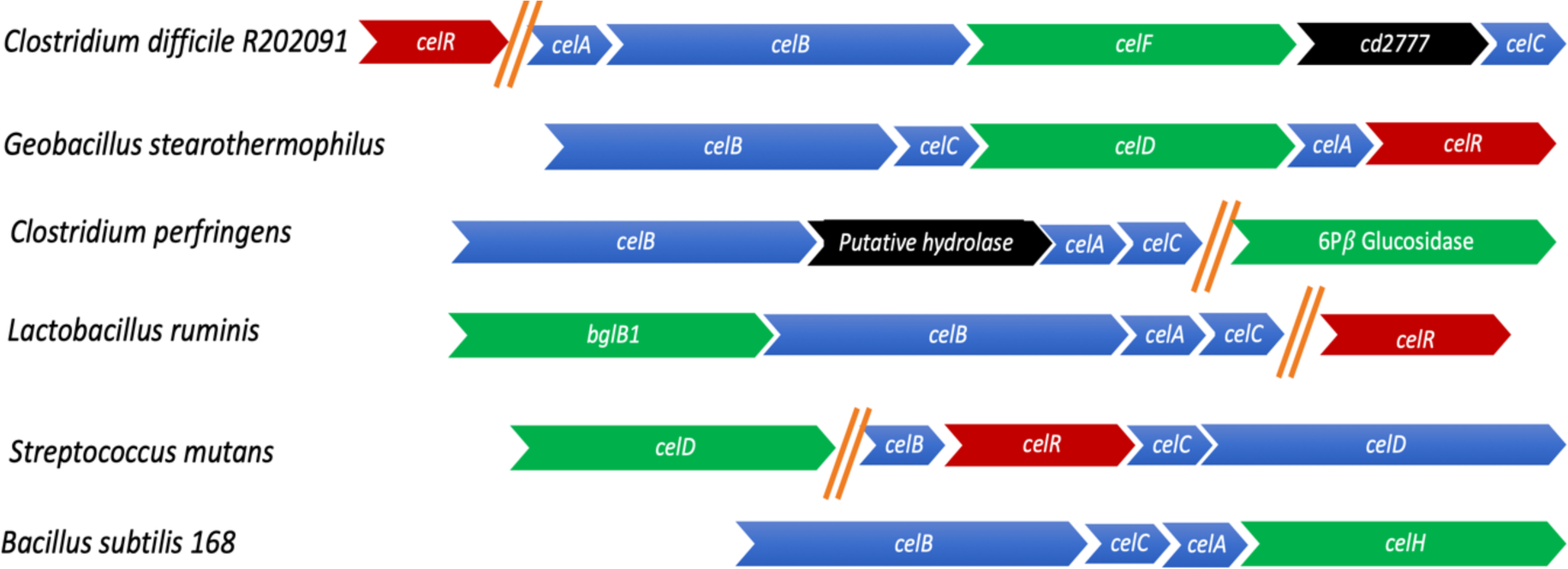
Cellobiose operon in selected Gram- positive bacteria. PTS components are colored in Blue. The Beta glucosidase is in green. Regulatory gene is colored in Red. Any additional genes are colored in Black. Double slash indicates genes are not in the same operon and are transcribed separately.

**Figure S2.**
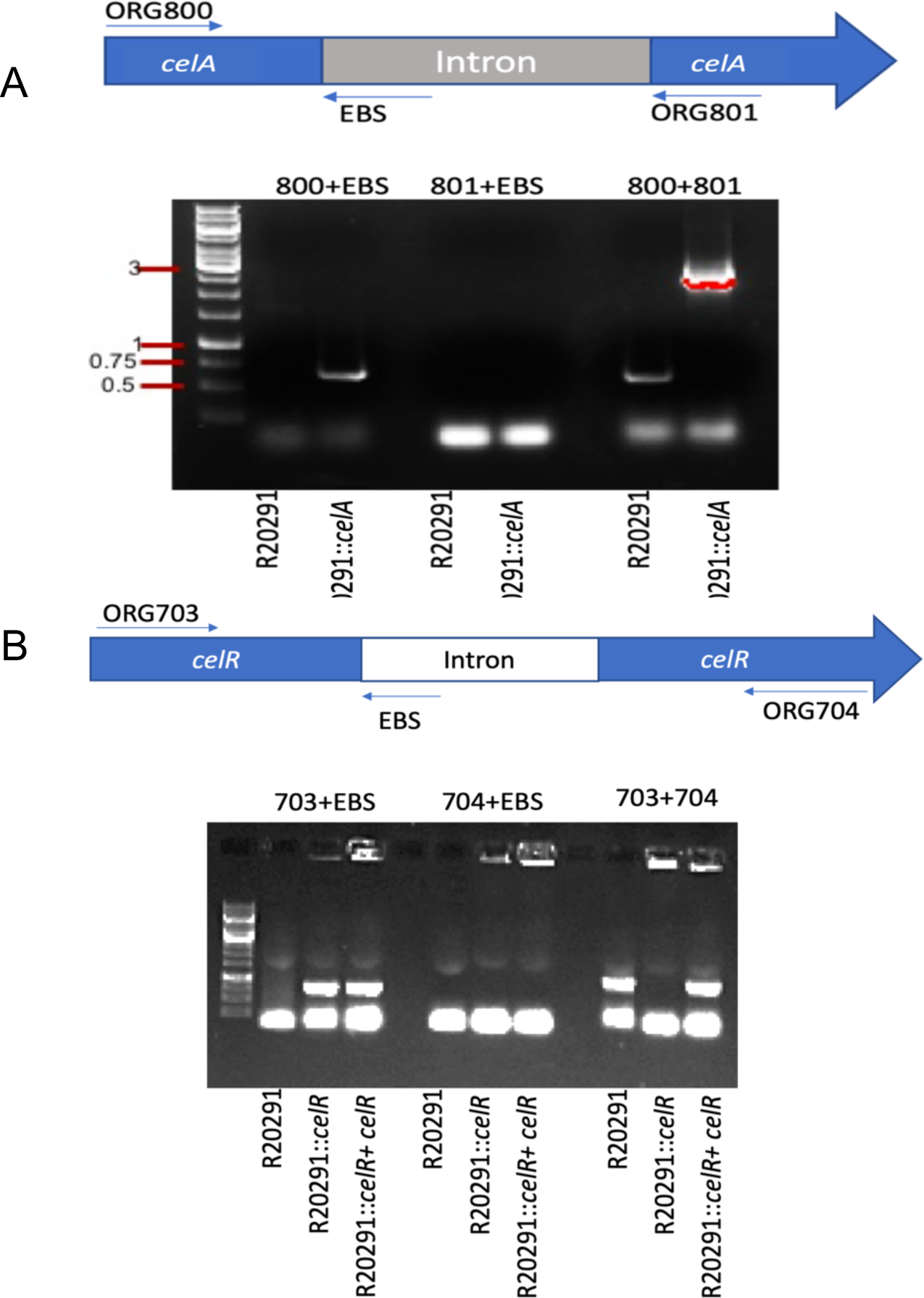
Construction and confirmation of the lic operon mutant in *C. difficile* R20291strain. **A.** ClostTron (group II intron) mediated insertional inactivation of *celA* the first gene in the *lic* operon. PCR verification of the intron insertion in *celA in* R20291strain, conducted with intron-specific primer EBS universal [EBS(U)] with *celA* specific primers ORG800 and ORG801. **B.** ClostTron (group II intron) mediated insertional inactivation of *celR*. PCR verification of the intron insertion in *celR in* R20291strain, conducted with intron-specific primer EBS universal [EBS(U)] with *celA* specific primers ORG703 and ORG704. Mutant complemented with celR (R20291::*celR*+*celR*) is also included in the analysis.

**Figure S3.**
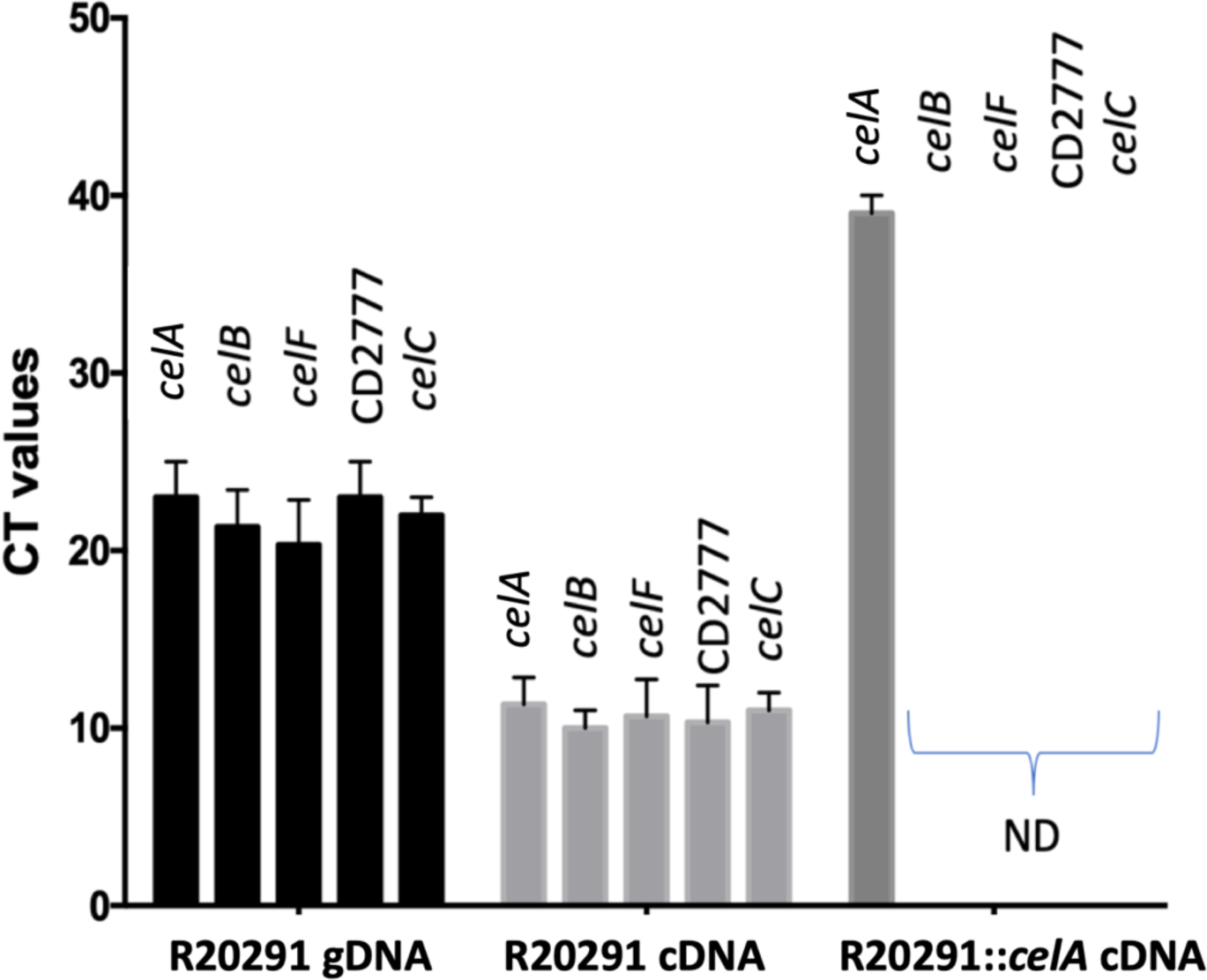
qRT-PCR to detect transcripts of genes in *lic* operon. R20291 genomic DNA (10ng) was used as a positive control. cDNA was prepared from the total RNA was harvested from 8h old R20291 and R20291::*celA* mutant cultures. Higher the CT value indicates lower transcripts level; ND (Not Detectable).

**Figure S4.**
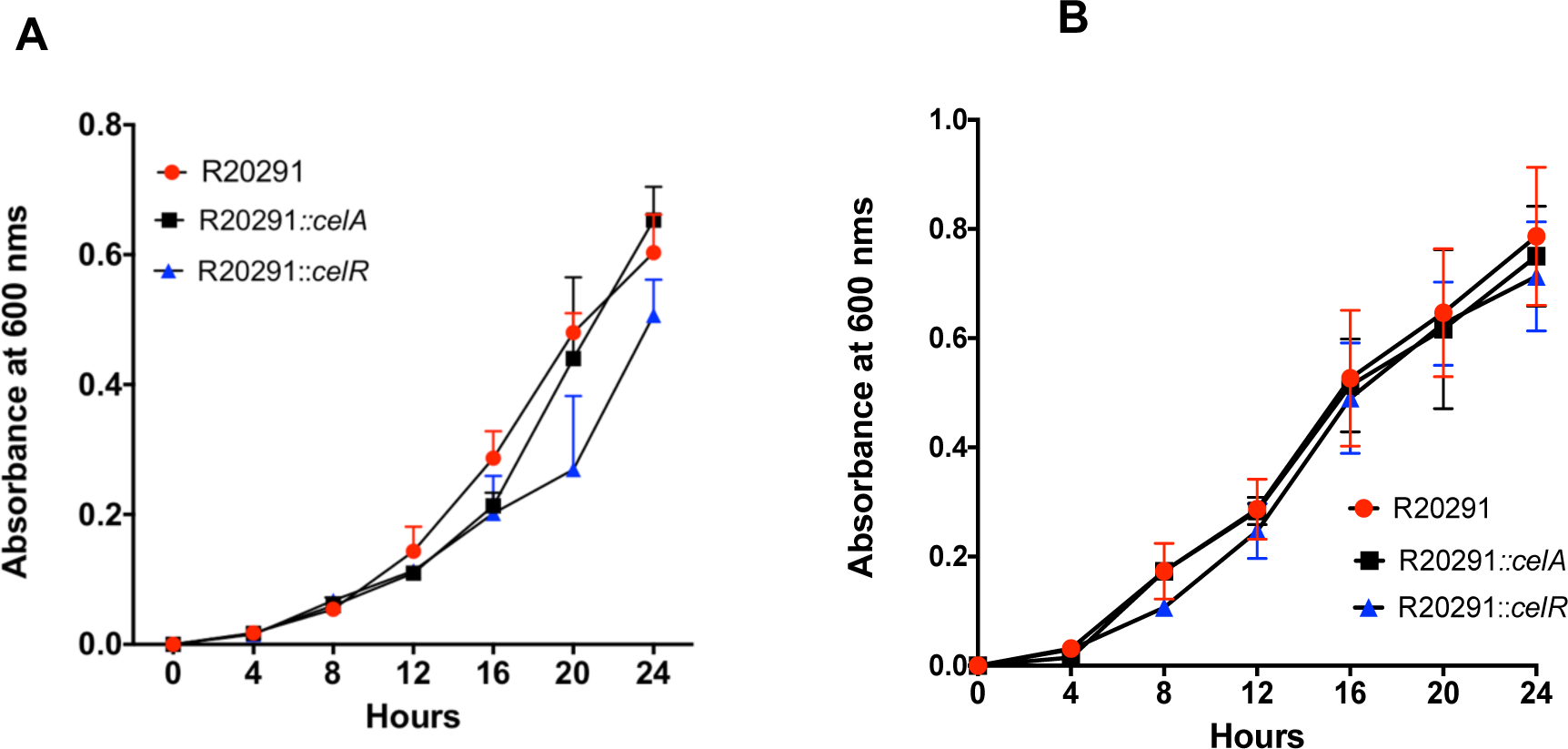
Growth comparison between R20291, R20291::*celR*, and R20291::*celA* strains grown in minimal media supplemented with 0.1% chitibiose **(A)** or 20 mM N-acetylglucosamine **(B)**.

**Figure S5.**
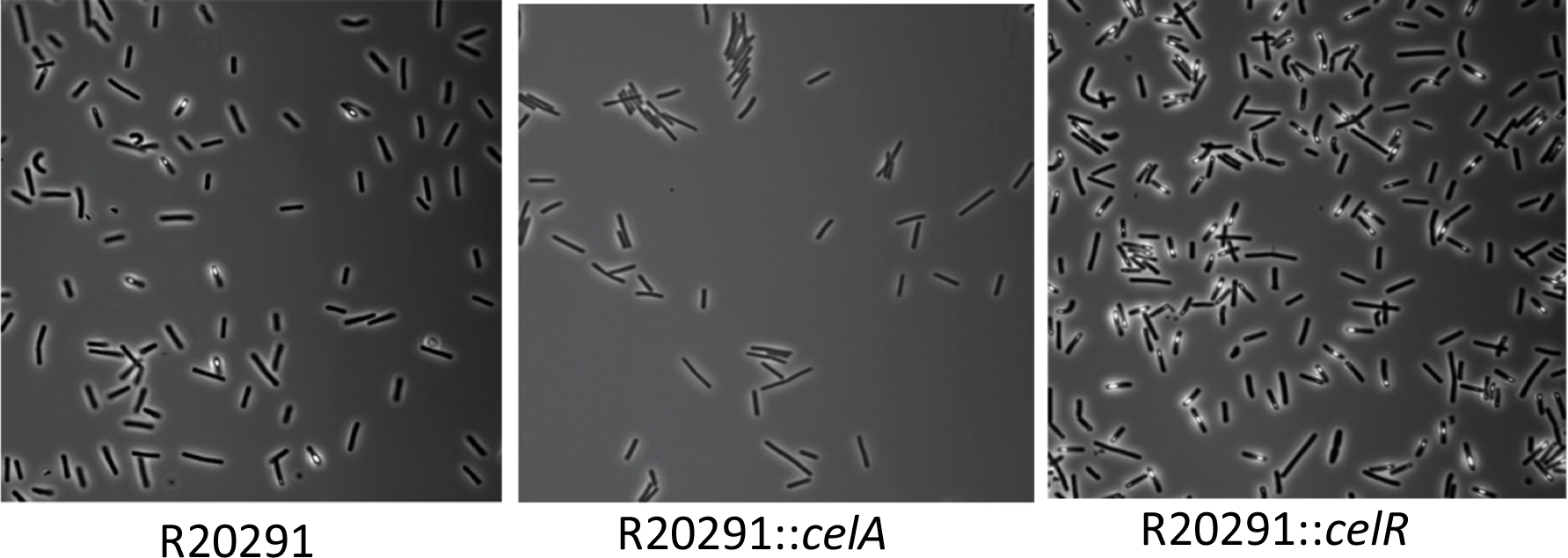
Microscopic analysis of *C. difficile* strains for sporulation. Strains were grown in TY-agar for 36 hours before analysis. Spores are visible as while halo within the cells.

**Figure S6.**
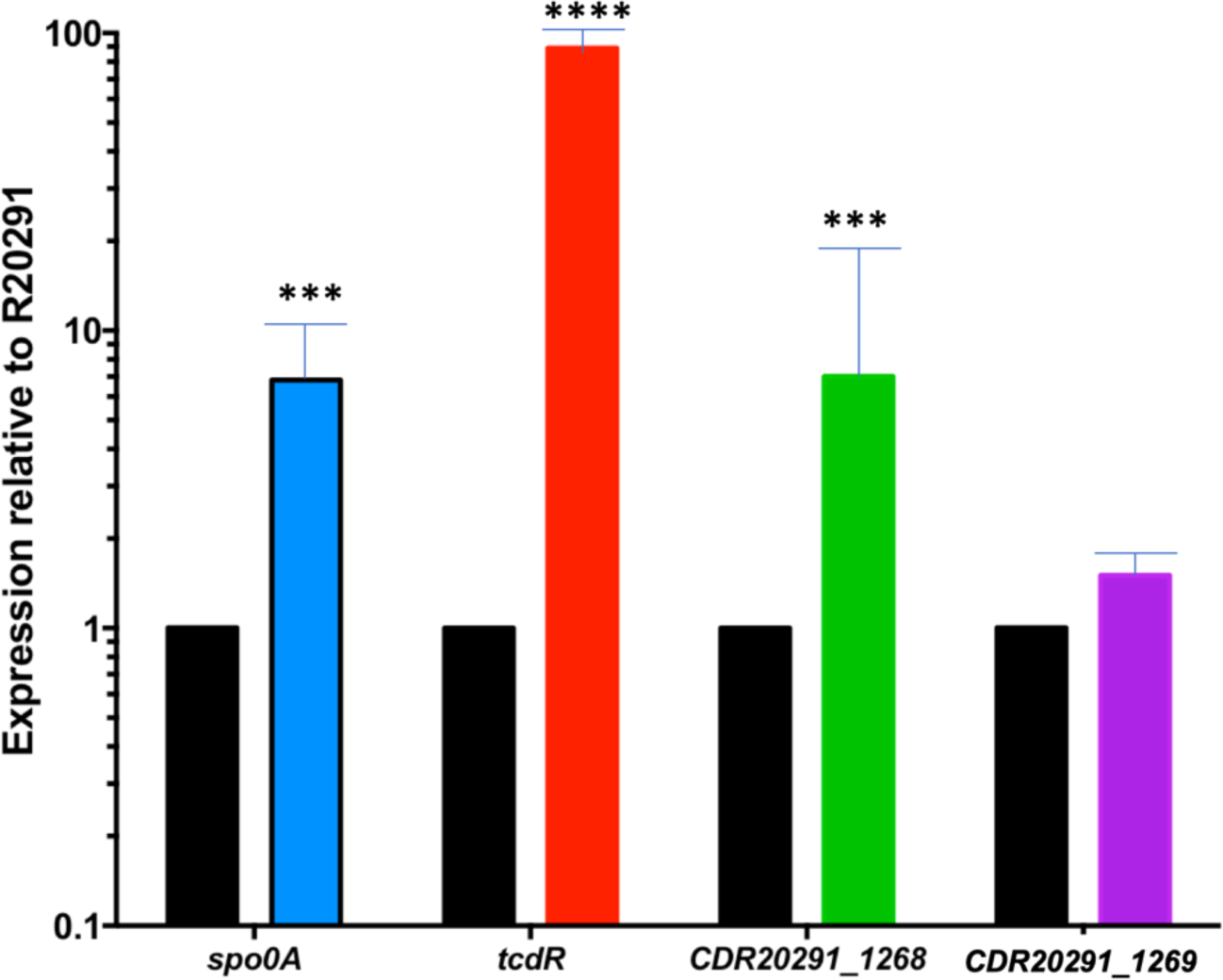
qRT- PCR to quantify *spo0A* and *tcdR* transcripts. Results of the R20291::*celR* for *spo0A* (sporulation regulator), and *tcdR* (regulator for toxin production) indicates a general trend of overexpression of these genes corroborating with the observed phenotypes of higher toxin production and sporulation in R20291::*celR* strain. ****=P<0.0001, *** P<0.001 using Student’s t-test.

**Table S1:** Bacterial strains and plasmids used in this study.

| Bacterial strain or plasmid | Relevant features or genotype | Source or reference |
| --- | --- | --- |
| *C. difficile* R20291 | Clinical isolate - NAP1/027 ribotype, isolated in 2006 following an outbreak in Stoke Mandeville Hospital, UK | Stabler *et al.* (2009) |
| *Escherichia coli* DH5α | *endA1 recA1 deoR hsdR17 (r<sub>K</sub><sup>-</sup> m<sub>K</sub><sup>+</sup>)* | NEB |
| *E. coli* S17-1 | Strain with integrated RP4 conjugation transfer function; favors conjugation between *E. coli* and *C. difficile* | Teng *et al.* (1998) |
| pMTL007-CE5 | ClosTron plasmid | Heap *et al.* (2010) |
| pMTL007-CE5:Cdi-*celA*-178-179a | pMTL007-CE5 with group II intron targeted to first gene (*celA*) of cellobiose operon | This study |
| pMTL007-CE5:Cdi-*celR*-228-229a | pMTL007-CE5 with group II intron targeted to *celR*, the putative transcriptional regulator of the cellobiose operon | This study |
| pRPF185 | *E. coli* / *C. difficile* shuttle plasmid | Fagan *et al.* (2011) |
| pRG381 | pRPF185 with *celR*-FLAG with 200 bps upstream DNA | This study |
| pRG382 | pRPF185 containing 200 bps upstream DNA with *PcelA-gusA* | This study |
| pRG383 | pRPF185 containing 600 bps upstream of CDR20291_2782 and *celR-gusA* | This study |
| pRG384 | pRPF185 containing 98 bps upstream of *celR* with *gusA* | This study |

**Table S2:**
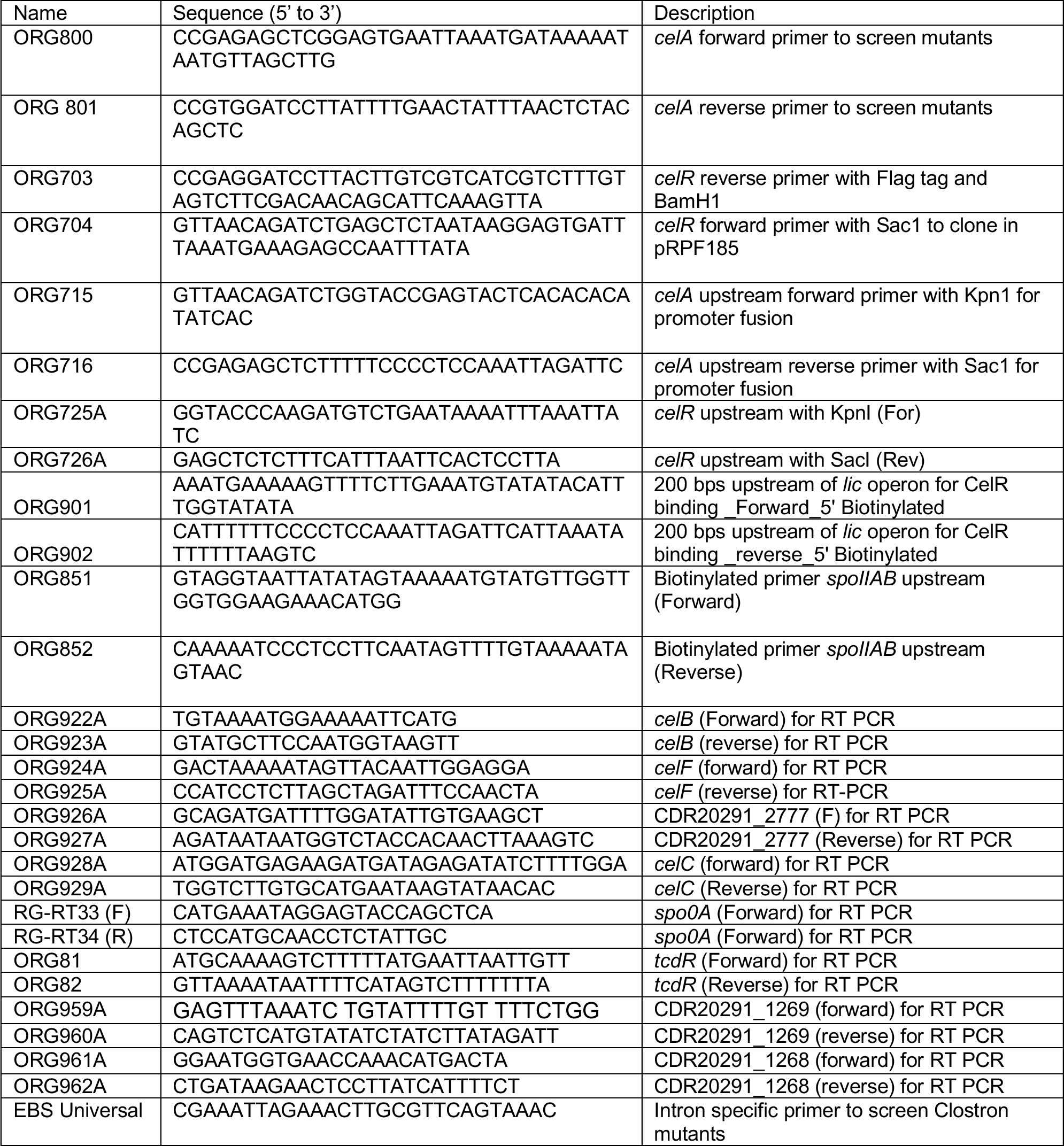
Oligonucleotides used in the study.

**Table S3.**
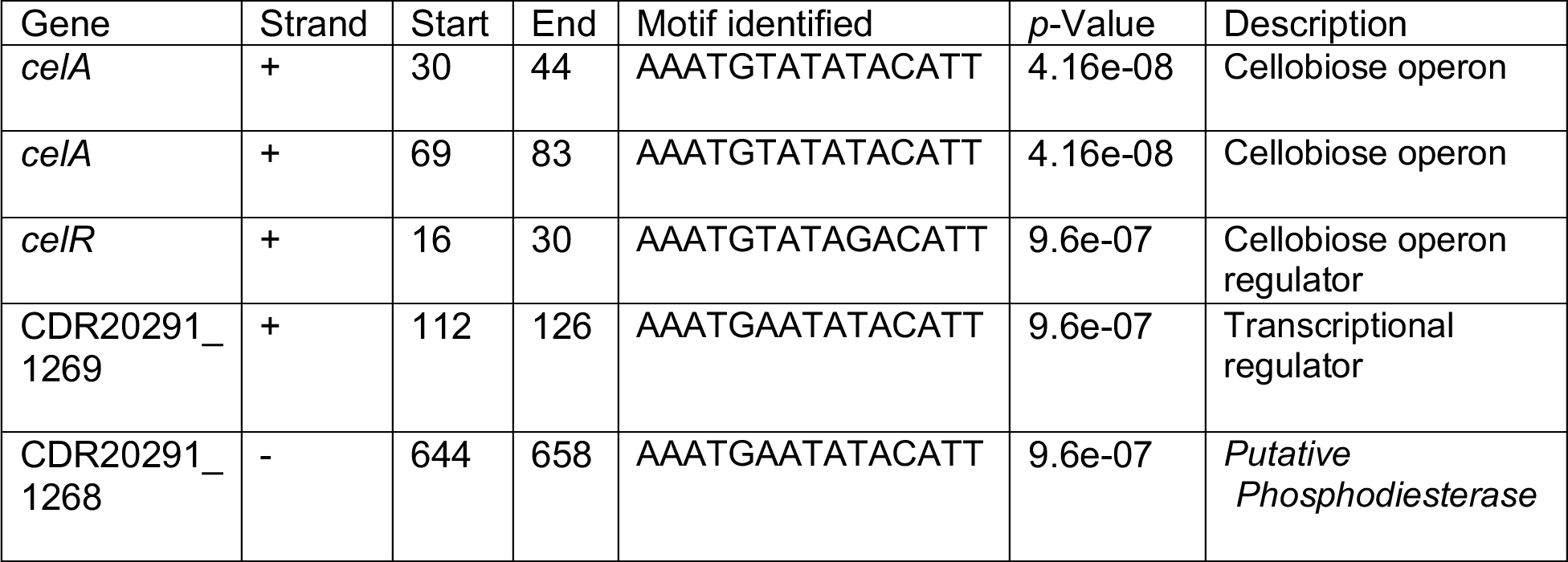
*C. difficile* R20291 genes with CelR binding motifs in their upstream region identified by FIMO motif search program.

